# Tracing 600 years of long-distance Atlantic cod trade in medieval and post-medieval Oslo using stable isotopes and ancient DNA

**DOI:** 10.1101/2024.01.25.577044

**Authors:** Lourdes Martínez-García, Angélica Pulido, Giada Ferrari, Anne Karin Hufthammer, Marianne Vedeler, Alex Hirons, Catherine Kneale, James H. Barrett, Bastiaan Star

## Abstract

Marine resources have been important for the survival and economic development of coastal human communities across northern Europe for centuries. Knowledge of the origin of such historic resources can provide key insights into fishing practices and the spatial extent of trade networks. Here, we combine ancient DNA and stable isotopes (δ^13^C, δ^15^N, non-exchangeable δ^2^H and δ^34^S) to investigate the geographical origin of archaeological cod remains in Oslo from the eleventh to seventeenth centuries CE. Our findings provide genetic evidence that Atlantic cod was obtained from different sources, including a variety of distant-water populations like northern Norway and possibly Iceland. Evidence for such long-distance cod trade is already observed from the eleventh century, contrasting with archaeological and historical evidence from Britain and other areas of Continental Europe around the North and Baltic Seas, where such trade developed in a later period. Diverse biological origins are further supported by significant differences of a range of isotopes, indicating that multiple populations living in different environments were exploited. This research highlights the utility of combining ancient DNA methods and stable isotope analysis to describe the development of marine fisheries during the medieval and post-medieval period.

## 1. Introduction

Understanding the extent of historic Atlantic fisheries reveals the importance of marine resource exploitation for medieval and post-medieval European societies [1]. For instance, long-distance fish trade from production sites in the north (e.g., northern Norway and/or Iceland) to urban centres in Britain and mainland Europe is well-documented by historical and archaeological sources for medieval and early modern times, being especially evident by the thirteenth and fourteenth centuries [2–4]. Nonetheless, for the earlier Middle Ages, temporal and spatial patterns of exploitation and long-distance trade remain poorly documented. While the earliest known example of long-distance trade of Atlantic cod (*Gadus morhua*) presently has a *terminus ante quem* of *ca.* 1066 CE (by which date northern Norwegian cod was brought to Haithabu in what is now Schleswig-Holstein) [5], the species was predominantly locally acquired in England and Flanders during the tenth to twelfth centuries [6–8]. Thereafter, an increasingly commercialized long-range trade of air-dried Atlantic cod (*stockfish*) only appeared from the thirteenth to fourteenth century onwards, around the southern North Sea and the eastern Baltic Sea [6–8]. This dried fish was likely traded via Bergen to medieval centres across Europe (Germany, Sweden, Poland, Estonia, England) [3, 6, 7, 9–12]. Nonetheless, the development of the early medieval Atlantic cod trade remains to be discovered, between the early outlier of *ca.* 1066 CE at Haithabu, and the more widespread boom of the thirteenth and fourteenth centuries. A promising location to do so is within the milieu of medieval Scandinavia, where processed fish, in particular *stockfish* was produced and found a ready cultural reception within local foodways [13]. For instance, Oslo, in Norway, emerged as a town during the tenth to eleventh centuries [14, 15]. Large numbers of fish remains (especially of Atlantic cod) [4] have been found during archaeological excavation of Oslo’s urban settlement layers [16, 17]. By the fourteenth century, Oslo had become an important town and centre of consumption, although not a major hub for the transhipment of processed fish. By that time, participation in long-range fish-trade is likely to be present. However, earlier patterns, and trends of trade through time, remain poorly understood.

Recent advances in ancient DNA (aDNA) approaches have provided insights about the usage of fish across time [18], its historical declines [19] and distributional fluctuations [20], and offer opportunities to investigate the biological origin of archaeological fish bones [5, 8]. Similarly, stable isotopes represent a complementary method to aDNA for such purposes, as isotopes can record information about diet and habitat use and/or environmental conditions of different species, including fish [6, 21–23]. Therefore, determining the biological origin of economically important species like Atlantic cod using the combined inference of aDNA and stable isotope methods can provide novel evidence of the expansion of medieval marine fisheries and explores the potential increase in the exploitation of targeted populations, beyond what is needed for local subsistence. Here, we use such a multidisciplinary approach, combining genome-wide aDNA and stable isotopes to describe the development of long-distance Norwegian trade to Oslo, over a period of approximately 600 years during the medieval and post-medieval periods, starting *ca.* 1000 CE.

To investigate the biological origin of Atlantic cod from medieval Oslo, we analyzed a total of 106 archaeological specimens using low-coverage whole-genome aDNA approaches (50 out of the 106 specimens) and/or stable carbon (δ^13^C), nitrogen (δ^15^N), non-exchangable hydrogen (δ^2^H) and sulphur (δ^34^S) isotope values (100 out of the 106 specimens). We aim to identify biological differences within these specimens (given their isotopic signatures, bone element and estimates of fish size), and to determine their geographical source based on genome-wide sequencing. Integrating both approaches, our observations support a diverse origin of Atlantic cod specimens in this assemblage, including specimens obtained through long-distance trade since *ca.* 1000 CE.

## 2. Material and methods

### Sample collection

We analysed 106 Atlantic cod bone samples collected from two archaeological sites: Oslogate 6 (*n* = 100) and Oslo Mindets tomt (*n* = 6; Table S1). The zooarchaeological assemblages (bones) from both sites are stored in the osteological collections at the University Museum, University of Bergen. Oslo was one of the first Norwegian towns, founded in the tenth to eleventh centuries. Oslogate 6 (59.91°N – 10.77°E) was located in the northern part of medieval Oslo [17]. This site was excavated during 1987-1989, while Oslo Mindets tomt (59.90°N – 10.76°E) was excavated in 1973 (Figure 1). Findings in these archaeological sites highlighted agriculture, fishing and metalworking as common activities in the area [24–28]. Remains of barley, fish-bones and animal dung (likely from livestock) have been found in the area, while increases in large leather deposits by the late twelfth century in Oslogate 6 are indicative of the development of trading activities (i.e., shoemaking) in the town [17, 24].

**Figure 1.**
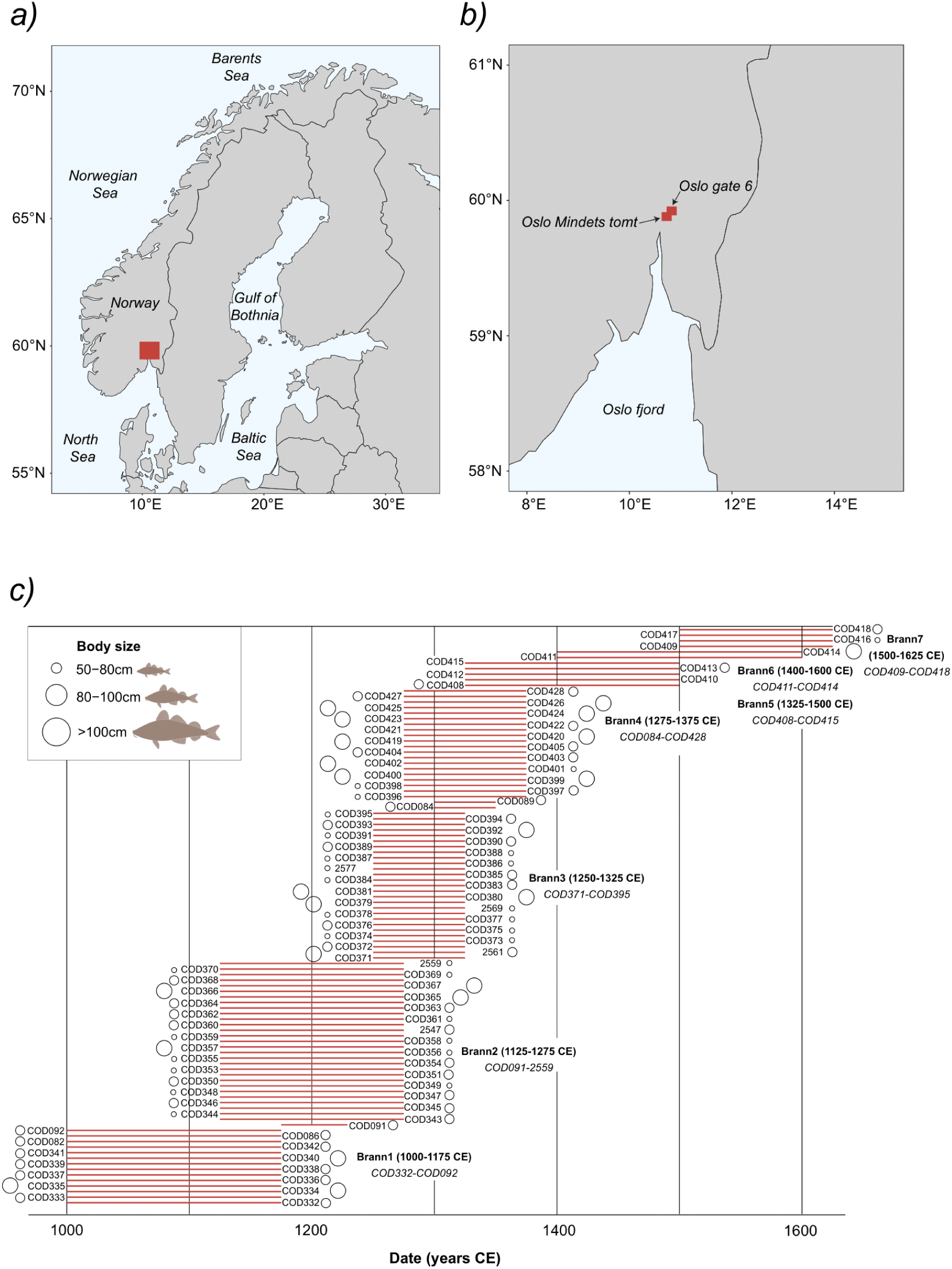
Geographical location of the archaeological Atlantic cod specimens collected from two archaeological sites in ***(a)*** southeast Norway in ***(b)*** Oslo (Oslogate 6, *n* = 100 and Oslo Mindets tomt, *n* = 6; Table S1). ***(c)*** Date distribution and body size estimation for archaeological samples. Specimens are obtained from layers placed between distinct fire layers (*Branntrinn*) that have been dated using archaeological methods (across the eleventh to seventeenth centuries). A total of eight fire stages (Brann1 to Brann8) have been identified and are used to describe the stratigraphy and constructions in the urban development [17]. Specimens COD406 and COD407 (from Brann8) do not have a confident date estimation and are excluded from this figure. Body size is significantly different between fire layers (Kruskal- Wallis chi-squared = 11.93, df = 3, *p-value* = <0.01), with fish from Brann4 significantly larger than those from Brann2 (TukeyHSD post-hoc test *p*-value = 0.04) and Brann3 (TukeyHSD post-hoc test *p-value* = 0.04).

Samples –cranial and postcranial elements– were stored dry and unfrozen after field collection. All specimens were morphologically and genetically identified as Atlantic cod. Samples are dated based on archaeological context (e.g., fire layer). Specifically, samples at Oslogate 6 are distributed between distinct fire layers (*Branntrinn*) that have been dated using archaeological methods (across the eleventh to seventeenth centuries). A total of 7 main fire stages (Brann1- 7) have been found in the excavation area and are used to described the stratigraphy and constructions in the urban development (i.e., waste layers) [for details about the fire layers see 17]. An eighth-layer (Brann 8) has been described for Oslogate 6, which includes an uppermost layer that is associated with modern times. Samples across the fire layers at Oslogate 6 are distributed as follows (Table S1): Brann1 (oldest) (where 12 out of 14 samples were processed for genomic analysis), Brann2 (*n* = 7/30), Brann3 (*n* = 8/27), Brann4 (*n* = 10/22), Brann5 (*n* = 5/5), Brann6 (*n* = 2/2), Brann7 (*n* = 4/4), and Brann8 (n = 2/2). Oslo Mindets tomt samples are dated by archaeological context [26] and are analysed together with those specimens from Oslogate 6 with which they co-occur in time. Three specimens (COD082, COD086, COD092) co-occur in time with Brann1, one specimen (COD091) co-occurs with Brann2, and two specimens (COD084, COD089) co-occur with Brann 4.

### aDNA extraction, library preparation and sequencing

A total of 50 samples were processed in the aDNA laboratory at the University of Oslo [29, 30] (Table S1, S2). Samples were treated as per Ferrari, Cuevas [31] and Martínez-García, Ferrari [32] before DNA extraction. In short, fish-bones were UV-treated for 10 minutes per side and milled with a stainless-steel mortar [33]. Starting material for DNA extraction consisted of two aliquots of fish-bone powder per specimen (150-200 mg per aliquot). Genomic DNA was extracted using a mild bleach treatment and pre-digestion step (BleDD) protocol [34]. Double-indexed blunt-end sequencing libraries were built from 15 or 20 μl of DNA extract using either the Meyer-Kircher protocol [35, 36] with modifications by Schroeder, Ávila-Arcos [37], or the single-stranded Santa Cruz Reaction (SCR) protocol (tier 4) [38] (Table S1). Library quality and concentration were examined with a High Sensitivity NGS Fragment Analysis Kit on the Fragment AnalyzerTM (Advanced Analytical). Libraries were sequenced on the Illumina HiSeq 4000 or on the Novaseq 6000 platform at the Norwegian Sequencing Centre (Table S1) and demultiplexed allowing zero mismatches in the index tag. Sequencing reads were processed using PALEOMIX v1.2.13 [39] and AdapterRemoval v.2.1.7 [40] to trim residual adapter contamination, filter and collapse of reads. Filtered reads were aligned using the gadMor2 genome as reference [41, 42] using BWA v.0.7.12 [43] with the *backtrack* algorithm, disabled seeding and minimum quality score of 25. aDNA deamination patterns were characterized using MapDamage v.2.0.9 [44].

### Genomic analysis

To determine the biological origin of 50 Atlantic cod specimens, we followed the hierarchical approach described in Martínez-García, Ferrari [8]. First, we used the genome-wide approach in the BAMscorer pipeline [45] to assign ancient cod specimens to the eastern Atlantic or the western Atlantic Ocean. Second, amongst those specimens with an eastern Atlantic origin, we use a similar approach to identify any specimen with a Baltic Sea origin. These assignments can be performed with high confidence (100% probability), given minimal data requirements (i.e, *ca.* 1000 reads that map towards the reference genome). Third, we used the chromosomal inversion approach in the BAMscorer pipeline [45] to determine the individual haplotypes of the four major chromosomal inversions in Atlantic cod (LG1, LG2, LG7 and LG12) [46]. The chromosomal inversions in Atlantic cod are associated with migratory behaviour and temperature clines [46–50]. Therefore, their combined genotype distributions can indicate their affinity towards a particular ecotype [5, 8]. Specifically, the probability of obtaining an inversion genotype follows a binomial distribution given the underlying allele frequency in a population. It is therefore possible to calculate the overall probability of obtaining a composite ancient inversion genotype —based on modern populations’ respective allele frequencies— as a measure of an individual’s affinity toward a specific population [5]. We included comparative (modern) inversion frequencies from the following populations: the Northeast Arctic (NEA), Iceland (frontal and coastal ecotypes, which differ in their tendency for long-distance migration), Norwegian Coast (Lofoten and southwest), the North Sea, the Irish Sea and Øresund (Figure 2a) [5, 50, 51]. We consider the possible uncertainty associated with sampling fixed alleles by re-calculating those allele frequencies where 1.0 and 0.0 frequencies were given to any of the chromosomal inversion genotypes. Thus, following the approach in the BAMscorer pipeline [45], we calculated a minimum expected frequency of (1/((2\**N*)+1)) for the minor allele, where *N* is the number of individuals in any population sample where alleles were fixed [5, 50, 51]. Samples with >0.1% endogenous DNA and an average of >50 single nucleotide polymorphisms (SNPs) across all four chromosomal inversions have been included for specific genomic assignments (Table S1). For each individual specimen, the source population with the highest percentage (%) has been considered as its most likely biological origin, although such assignment probabilities may remain low overall (Table S2, S3). Specimens with equal or similar assignment probabilities for two populations (e.g., 33% and 33%), have both populations as their putative origin (Table S2, S3). Finally, we recognized two spatially distinct groups (northernmost and north-central) as described in Martínez-García, Ferrari [8], that can be assigned with high and near bimodal probability. These groups are associated with population structure on a spatially larger scale (i.e., region). The northernmost group included NEA and Iceland (adding the probabilities of both Icelandic ecotypes), while the north-central group included the Norwegian Coast (coastal Atlantic cod from Lofoten and southwest Norway), the North Sea, the Irish Sea and Øresund [8]. While our ability to obtain high affinities of ancient specimens to specific populations within these two distinct regions is often low, we consider those that fall within the northernmost group (NEA and Iceland) with high probability (see below) as specimens that must have been obtained through long-distance trade following Martínez-García, Ferrari [8]. In contrast, we cannot exclude a putatively local biological origin (i.e., near Oslo) for those specimens belonging to the north-central group.

**Figure 2.**
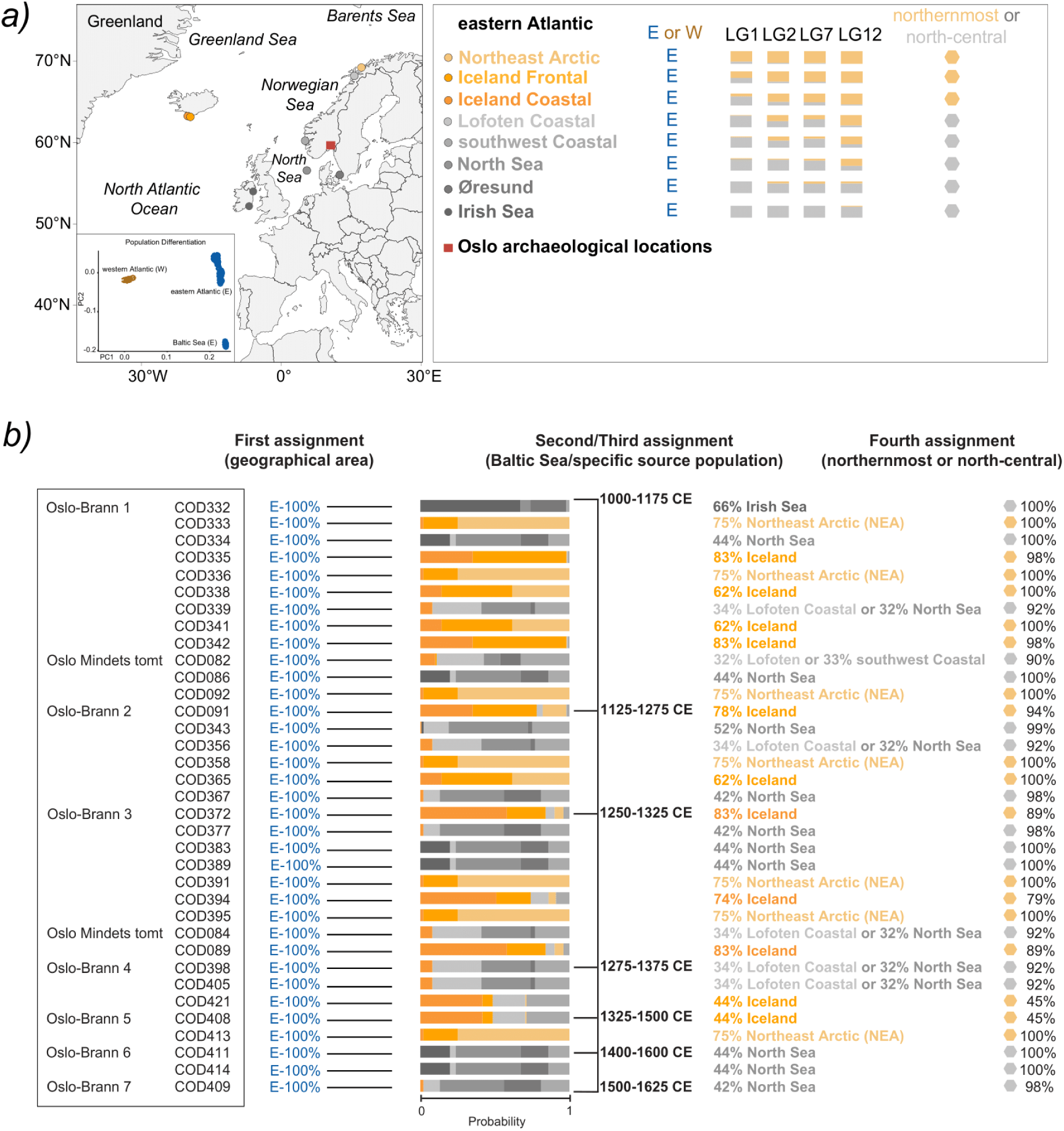
Genetic analyses of archaeological Atlantic cod specimens from Oslo. ***(a)*** Geographical distribution of modern inversion frequencies of chromosomal inversions in Atlantic cod (LG1, LG2, LG7 and LG12) from reference populations across the North Atlantic Ocean. Map is modified from Martínez-García, Ferrari [8]. Alleles associated with a northernmost (Northeast Arctic and Iceland) composite genotype distribution are assigned in orange. Alleles associated with a north-central (Norwegian Coast, the North Sea, the Irish Sea, Øresund) genotype distribution are assigned in grey. Oslo’s archaeological sites are indicated with a red square. ***(b)*** Genomic assignment to a geographical area, a source population and a genotypic group is based on the frequencies of chromosomal inversions of Atlantic cod (LG1, LG2, LG7 and LG12) as per Star, Boessenkool [5]. Percentages (%) indicate the highest probability to be from one population and area.

Furthermore, we investigated possible associations between cranial [premaxilla, articular, dentary, maxilla] or postcranial [vertebra and cleithrum]) bone elements and the putatively local (north-central group) or long-distance (northernmost group) origin of the specimen using a Fisher’s exact test. The test was implemented using the *stats* and *ggstatsplot* packages in R [52] for 27 specimens that have been assigned with a >70% probability to a northernmost or north-central genotype composite (Table S2). We excluded samples that have not been confidently assigned to a source population (COD082, COD084, COD339, COD356, COD398, COD405) and samples with an indistinguishable origin (low probability assignment to a northernmost or north-central genotype composite; COD408 and COD421) as per Martínez-García, Ferrari [8] (Table S2).

Additionally, we investigated if the proportion of specimens coming putatively either from the north-central or the northernmost group, and northern Norway or Iceland as individual locations (both locations were centres for *stockfish* production) changed over time. We used a Fisher’s exact test to evaluate the binary genetic assignment to the north-central or northernmost groups across time (>70% probability); while we evaluated if the probability – scaled from 0 to 1– of having a Northeast Arctic (NEA) or Icelandic origin (adding the probabilities of both Icelandic ecotypes) differs across time (i.e., fire layers), body size or bone element. Here, we used logistically transformed probabilities (*logit*) of NEA or Icelandic origin after testing for normality with a Shapiro-Wilk test. Thereafter, we performed non-parametric Kruskal-Wallis tests between such variables. Only fire layers with sufficient sample sizes (*n* ≥ 4 in Brann1 (1000-1175 CE) to Brann4 (1275-1375 CE)) were used in these analyses (*n* = 29). We excluded samples with a low probability assignment to a northernmost or north-central genotype composite (i.e., COD421). Finally, we investigated if there were significant differences between the size categories (50-80 cm, 80-100 cm and >100 cm) of specimens genetically assigned to different groups or populations (either from the north-central group, the northernmost group, or northern Norway or Iceland as individual locations) using a Fisher’s exact test. Analyses were computed in R using the *stats, ggstatsplot* and *car* packages [52, 53].

### Stable isotope analysis

Stable isotope analysis was performed on 100 Atlantic cod specimens (50 of which were also processed for genomic sequencing as noted above; Table S1). We measured stable carbon (δ^13^C), nitrogen (δ^15^N), non-exchangeable hydrogen (δ^2^H) (hereafter described as δ^2^H for simplicity) and sulphur (δ^34^S) isotopes on purified bone collagen. For collagen extraction, we used a cross-section of the bone to obtain an approximate life-time average for the isotopic estimates and thus reduce the impact of changing trophic level associated with age [54], which is captured within the concentric growth increments of acellular fish bone. Between *ca.* 400 and 650 mg of bone material was processed following the protocols of Barrett, Khamaiko [55] and references therein, with the exclusion of a lipid removal step (given the negligible level of fat present in archaeological cod bones). For δ^13^C and δ^15^N, the extracted collagen was analysed in triplicate at the Godwin Laboratory, Department of Earth Sciences, University of Cambridge, using a Costech elemental analyser coupled to a Thermo Finnigan Delta V IRMS. For δ^34^S and δ^2^H, (measured by Iso-Analytical Limited, by elemental analyser isotope ratio mass spectrometry), analyses were routinely in duplicate, although some samples yielded only enough collagen for single measurements. All isotope data are reported using international scales: δ^13^C values are reported relative to VPDB, δ^15^N values to AIR, δ^2^H values to VSMOW and δ^34^S to VCDT. The reported non-exchangeable δ^2^H values are corrected for exchangeable hydrogen by three-point linear calibration using standards. Overall, 64 samples passed appropriate quality-control thresholds for all four measured isotope ratios (atomic C/N ratio of 2.9 – 3.6, atomic C/S ratio of 125 – 225, atomic N/S ratio of 40 – 80) [56]. An additional 22 samples only passed appropriate C/N ratio quality-control thresholds. Therefore, the δ^13^C, δ^15^N and δ^2^H data are likely reliable for these 22 specimens, but potentially not the δ^34^S values. For this reason, analyses using only δ^13^C, δ^15^N and/or δ^2^H values (individually and interacting) are based on 86 specimens, while analyses including δ^34^S values (individually and in combination with other data) are based on 64 specimens.

We investigated correlations between C:N ratios and δ^13^C values, and correlations between δ^13^C-δ^15^N, δ^15^N-δ^2^H and δ^13^C-δ^34^S values after evaluating the normality of our data with a Shapiro-Wilk test using the *base* package in R [52]. We implemented a Spearman’s correlation (not normal distributed data) between C:N ratios and δ^13^C values (*n* = 86), and between δ^13^C- δ^34^S (*n* = 64); while Pearson’s correlations (normal distributed data) were used between δ^13^C- δ^15^N (*n* = 86) and δ^15^N- δ^2^H (*n* = 86). Correlations were computed in R using the *stats* package. Considering the environmental and ecological shifts that can occur through time and based on age and growth, we investigated variability of isotope values across time and across body size groups. We performed ANOVA analysis for δ^13^C, δ^15^N (*n* = 86) and δ^2^H values (*n* = 83, excluding individuals with missing δ^2^H values: COD407, COD419 and COD428) followed by a TukeyHSD post-hoc test; while a non-parametric Kruskal-Wallis test was computed for δ^34^S (*n* = 64) followed by a Dunn post-hoc test with a Bonferroni correction (<3 groups).

To identify potentially different ecological groups within Atlantic cod individuals, we examined the individual distribution of isotope values (δ^13^C, δ^15^N and δ^2^H) within fish across the three body size categories visualizing a density plot in R. We further evaluated the differences between the body size of all archaeological specimens distributed across Brann1 to Brann4 (*n* = 86) using a Kruskal-Wallis test, followed by a post-hoc Dunn test with a Holm correction (>3 groups). All analysis were performed in R using the *stats*, *FSA* and *ggplot2* packages [52, 57, 58].

### Combined genome-isotope analyses

A summary of isotope values, in relation to body size (50-80 cm, 80-100 cm and >100 cm) and the putatively genomic assignment of individuals to either a NEA or Icelandic origin, or a north-central or northernmost origin was represented by a principal component analysis (PCA) on the 64 relevant δ^13^C, δ^15^N, δ^2^H and δ^34^S individual estimates. We performed ANOVA analysis to describe the contribution of size and time (Brann1 to Brann4) to each principal component (PCs) in the PCA followed by a TukeyHSD post-hoc test. PCAs were plotted in R using the *ggplot2* package [58]. Loadings, eigen values, and contributions of each isotope to the principal components (PCs) of the PCA were calculated in R using the *factoextra* package [59].

To investigate if the (genomic) probability –scaled from 0 to 1– of having a NEA or Icelandic (adding the probabilities of both Icelandic ecotypes) affinity has different isotopic signatures, we computed a multivariate linear regression with logistically transformed probabilities (*logit*) of NEA or Icelandic origin (*n* = 20) as dependent variables, and δ^13^C, δ^15^N, δ^2^H and δ^34^S values as predictors. We tested the regression for homoscedasticity with a Breusch-Pagan test and validated the model by assessing residuals and symmetry. We further evaluated for multicollinearity by removing δ^15^N from our regressions. We did not observe major differences without δ^15^N; therefore, the final regression model includes all four isotope values. Additionally, considering that *stockfish* is primary produced from the Atlantic cod migratory ecotype in Norway, we selected those specimens with an Icelandic frontal (Icelandic migratory ecotype) affinity (another source of *stockfish* production), to assess the differences in isotopic signatures between migratory ecotypes. Multivariate linear regressions with specimens with an Icelandic frontal affinity follow the same procedure previously explained. All regressions and the Breusch-Pagan test were computed in R using the *stats* and the *car* packages [52, 53].

Finally, we assessed the direct relation between a binary assignment to NEA or Iceland (adding both ecotypes, migratory and stationary behaviour) and the δ^13^C, δ^15^N, δ^2^H and δ^34^S values. We used one-way ANOVAs (for δ^13^C, δ^15^N and δ^34^S) or Welch’s ANOVAs (for δ^2^H) after testing for equal variances with a Bartlett test (normal distributed data). Only specimens with a >70% probability NEA or overall >70% probability Icelandic assignment were included (Table S2). Therefore, 10 specimens were tested against δ^13^C, δ^15^N, and δ^2^H values and nine specimens were tested against δ^34^S values. Icelandic frontal (migratory ecotype) assignments are usually shared with an Icelandic coastal (stationary ecotype) assignment (e.g., 43% and 35% respectively; see Table S3). Consequently, an Icelandic frontal assignment alone cannot be binary assigned with >70% probability of origin to a population. Thus, we only included samples with a confident (>70% probability) overall Icelandic origin. ANOVAs and equal variance tests were computed using the *base* package in R.

## 3. Results

### Genomic analysis

We successfully sequenced 35 Atlantic cod specimens with a total of ∼770 million paired reads, of which ∼111 million reads aligned with a range of 0.1% to 47% of endogenous DNA content per specimen (Table S2). As expected, patterns of DNA fragmentation and deamination rates are consistent with those of authentic aDNA (Figure S1). We found that all 35 specimens can be assigned to an eastern Atlantic origin (100% assignment probability; Figure 2b and Table S1, S2) of which 19 specimens had the highest affinity to the north-central group (56-100% assignment probability). No specimens were identified with a Baltic Sea origin. Within the north-central group, specific population assignments are uncertain with 10 specimens putatively assigned to the North Sea (42-52% assignment probability), one specimen to the Irish Sea (66% assignment probability), five specimens to both the North Sea (32% assignment probability) and the Norwegian Coast (Lofoten, 34% assignment probability), one specimen to the Norwegian Coast (Lofoten and southwest, 32% and 33% assignment probability), and two specimens to the southwest coast of Norway (26% assignment probability; Figure 2b and Table S3). Sixteen specimens had the highest affinity to the northernmost group (79-100% assignment probability) of which seven specimens were assigned to the Northeast Arctic (75% assignment probability) and nine were likely assigned to Iceland (62-83% assignment probability, adding the probabilities of both migratory and stationary ecotypes; Figure 2b and Table S3).

While postcranial bones have been associated with long-distance sources (i.e., NEA and Iceland) [8], we did not find a statistically significant association between the bone element (cranial or postcranial) and specimens with a local (north-central group) or traded origin (northernmost group; *p-value* = 0.37; *n* = 27; Figure S2a, Table S2). In addition, we did not find statistically significant differences over time between specimens genetically assigned to either a north-central or northernmost group (*p-value* = 0.77; *n* = 29; Figure S2b, Table S1). We did not find any statistical difference between the probability of having a NEA or Icelandic origin in bone element (NEA: Kruskal-Wallis chi-squared = 2.28, df = 1, *p-value* = 0.13 and Iceland: Kruskal-Wallis chi-squared = 0.13, df = 1, *p-value* = 0.72; *n* = 33; Figure S3a, S3b), in body size (NEA: Kruskal-Wallis chi-squared = 2.00, df = 2, *p-value* = 0.37 and Iceland: Kruskal-Wallis chi-squared = 0.25, df = 2, *p-value* = 0.88; Figure S3c, S3d) or across time (Brann1 to Brann 4; NEA: Kruskal-Wallis chi-squared = 0.74, df = 3, *p-value* = 0.86 and Iceland: Kruskal-Wallis chi-squared = 0.04, df = 3, *p-value* = 0.99; Figure S3e, S3f). Furthermore, we did not find statistical differences between the size categories of specimens genetically assigned to Iceland and NEA (*p* = 0.09, *n* = 16), the north-central group and Iceland (*p* = 0.68, *n* = 25), the north-central group and NEA (p = 0.46, *n* = 23), the north-central group, NEA and Iceland (*p* = 0.26, *n* = 32), and the north-central group and northernmost group (*p* = 1.0, *n* = 32; Table S4).

### Isotope analysis

We successfully extracted collagen from 93 out of 100 Atlantic cod specimens, with 86 passing quality thresholds for δ^13^C, δ^15^N and δ^2^H (Table S1). For δ^34^S, 64 specimens passed quality control thresholds. Collagen yields of all 93 extractions ranged from 1.5% to 10.6%. Isotope values that passed initial quality control thresholds ranged from -15.9‰ to -11.9‰ (mean - 13.9‰for carbon (δ^13^C), from +12.7‰ to +16.8‰ (mean +14.9‰for nitrogen (δ^15^N), from - 13.8‰ to +38.1‰ (mean +12.9‰) for hydrogen (δ^2^H), and from +6.0‰ to +17.5‰ (mean +14.0‰for sulphur (δ^34^S) after the initial quality control (Table S5).

The C:N ratios for the included data ranged from 3.0 to 3.6 and were negatively related to δ^13^C values (Figure S4, Table S1). This correlation was weak and not significant (Spearman’s rho = -0.19, p-value = 0.07). Therefore, we assume that *post mortem* processes did not majorly impact our isotopic data [60]. Isotopes were strongly correlated with a significant relation between δ^15^N and δ^13^C values (Pearson’s correlation = 0.46, *p-value* = <0.01; Figure S5a), and between δ^2^H and δ^15^N values (Pearson’s correlation = 0.51, *p-value* = <0.01; Figure S5b). No significant relationship was observed between δ^13^C and δ^34^S values (Spearman’s correlation rho = 0.18, *p-value* = 0.15; Figure S5c).

Body size significantly influences three isotope values (Table S6a): δ^13^C (ANOVA: *F-value* = 6.70, *p-value* = <0.01, *n* = 86; Figure S6a), δ^15^N (ANOVA: *F-value = 5.47, p-value = <0.01*, *n* = 86; Figure S6b) and δ^2^H (ANOVA: *F-value* = 7.38*, p-value* = <0.01, *n* = 83; Figure S6c). Such significant differences in δ^13^C, δ^15^N and δ^2^H values were found between the smallest (50- 80 cm) compared to the largest (>100 cm) body size categories (TukeyHSD post-hoc test *p- value* = <0.01; Table S6a). Interestingly, we observed a bimodal distribution within medium sized fish (80-100cm) for δ^13^C and δ^15^N but not in δ^2^H or smaller and larger body size categories (Figure S7). Furthermore, we found significant differences of δ^13^C (ANOVA: *F- value* = 5.91, *p-value* = <0.01, *n* = 81; Figure S8a), δ^15^N values (ANOVA: *F-value* = 3.94, *p- value* = <0.01, *n* = 81; Figure S8b), and δ^2^H (ANOVA: *F-value* = 3.67, *p-value* = <0.01, *n* = 79; Figure S8c) across time (Brann1 to Brann4; Table S6b). Specifically, Brann4 (1275-1375 CE) had significantly higher isotope values (δ^13^C *p-value* = <0.01; δ^15^N *p-value* = 0.03 and δ^2^H *p-value* = 0.01) compared to Brann2 (1125-1275 CE). Brann4 also had significant higher δ^13^C values (*p-value* = <0.01) compared to Brann3 (1250-1325 CE) and significantly higher δ^15^N values (*p-value* = 0.01) compared to Brann1 (1000-1175 CE). No significant correlations were obtained between δ^34^S and body size (Kruskal-Wallis chi-squared = 0.11, df = 2, *p-value* = 0.95, *n* = 64; Figure S6d) or across time (Kruskal-Wallis chi-squared = 4.69, df = 3, *p-value* = 0.20, *n* = 60; Figure S8d). We found significant differences between the body size of our archaeological specimens across time (Kruskal-Wallis chi-squared = 11.93, df = 3, *p-value* = <0.01). These differences can be observed between larger fish from Brann4 against Brann2 (TukeyHSD post-hoc test *p-value* = 0.04), and Brann3 (TukeyHSD post-hoc test *p-value* = 0.04).

### Combined genome-isotope analyses

Principal component analysis (PCA) was employed to investigate the relationship between isotope data and genomic assignments while controlling for fish size. The first two principal component axes (PC1 and PC2) explained 99.18% of the observed variation (Figure 3a). Body sizes significantly changed across PC1 (ANOVA: *F-value* = 8.85, *p-value* = <0.01, *n* = 64), where PC1 values increased with body size (Figure 3a). Such differences were significantly found across PC1 between the smaller (50-80 cm) compared to the medium (80-100 cm; TukeyHSD post-hoc test *p-value* = <0.01) and larger (>100 cm; TukeyHSD post-hoc test *p- value* = <0.01) body size categories. We found that the highest contribution to the observed variation in PC1 was that of δ^2^H (PC1 contribution = 99.53%), which is to be expected given the association noted above between δ^2^H values and body size (Figure 3a, Table S5a, S5b). Importantly, no body size changes were observed across PC2 (ANOVA: *F-value* = 0.12, *p-value* = 0.89, *n* = 64, Figure 3). PC2 presented a negative relation with δ^34^S according to its loading values (Table S5a). We found that the highest contribution to the observed variation in PC2 were δ^34^S values (PC2 contribution = 98%, Table S5b). Furthermore, values in PC3 increased with δ^15^N and δ^13^C values (Table S5a, S5b), however, no particular body size changes were observed across PC3 (ANOVA: *F*-value = 2.25, *p*-value = 0.11, *n* = 64). In the PCAs of isotope values highlighting the genomic assignment individuals, the second and third principal component axes (PC2 and PC3) explained 4.5% of the observed variation (Figure 3b, 3c). The north-central vs. northernmost genomic groups are separated on PC3, whereas NEA vs. Iceland do no separate in on PC2 or PC3 (Figure 3b, 3c).

**Figure 3.**
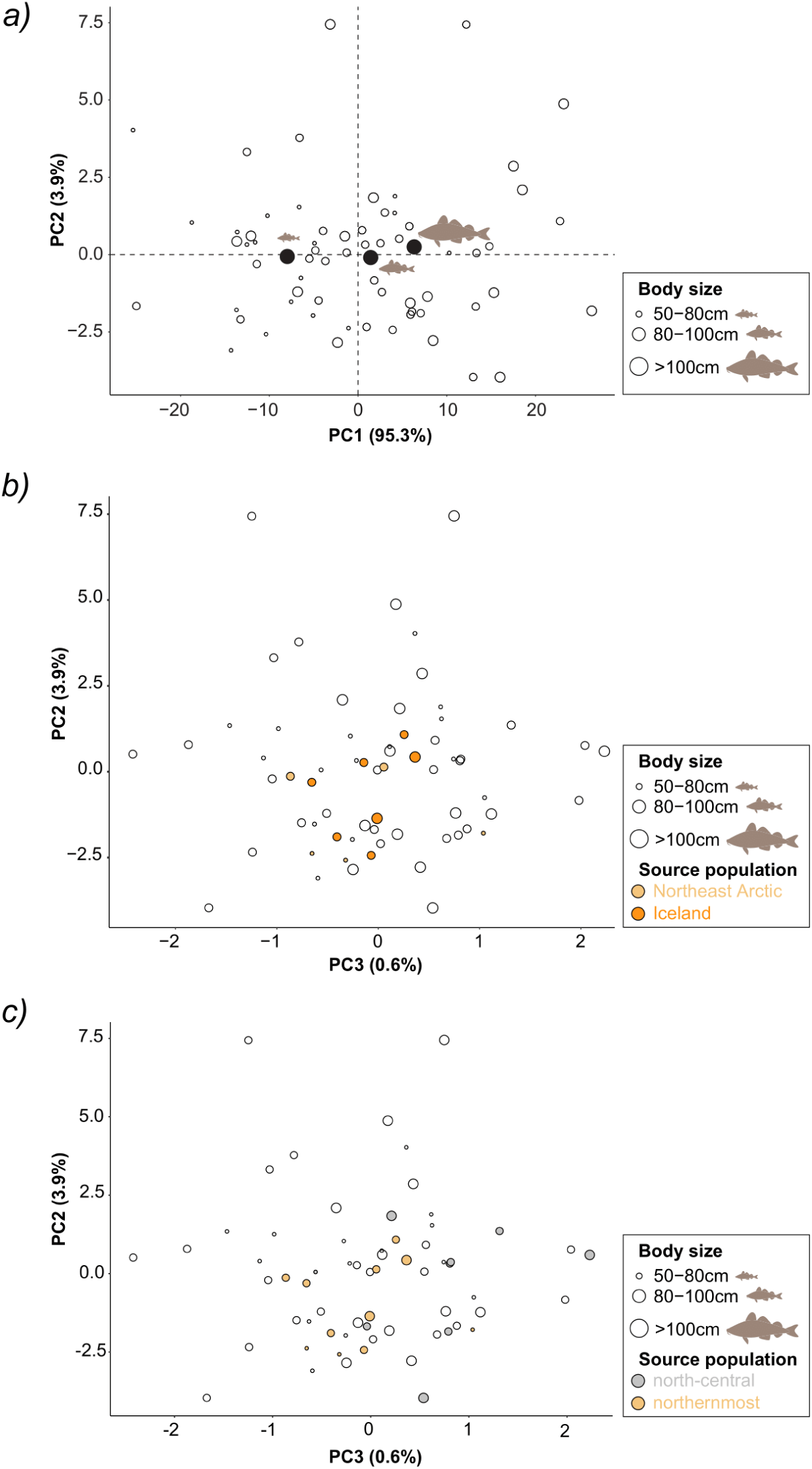
Principal component analysis (PCA) for carbon (δ^13^C), nitrogen (δ^15^N), non- exchangeable hydrogen (δ^2^H) and sulphur (δ^34^S) isotopes highlighting ***(a)*** body size and ***(b)*** putative genetically inferred source population assignments of individuals to the Northeast Arctic (NEA) or Iceland populations, and ***(c)*** a north-central or northernmost genomic groups. Values distributed across PC1-PC2 axes explain 99.18% of the observed variations, while values across PC2-PC3 axes explain 4.5% of the observed variation.

Multivariate linear regressions showed a significant relation between δ^13^C values and the (genomic) probability of having a northern Norway origin (NEA, *p-value* = 0.01, adjusted *R*- squared = 0.50; Table S7a), but not for the (genomic) probability of having an Icelandic (*p-value* = 0.36; Table S7b) or Icelandic frontal origin (*p-value* = 0.20; Table S7c). We found a significant difference between the δ^13^C values depending on the binary population assignment between a NEA or Icelandic biological origin (ANOVA: *F-value* = 5.99, *p-value* = 0.04, Figure 4). Such differences were not observed for δ^15^N, δ^2^H and δ^34^S (Figure S9a, S9b, S9c). Other isotope values did not show any significant influence on the biological origin of Atlantic cod specimens (Table S7).

**Figure 4.**
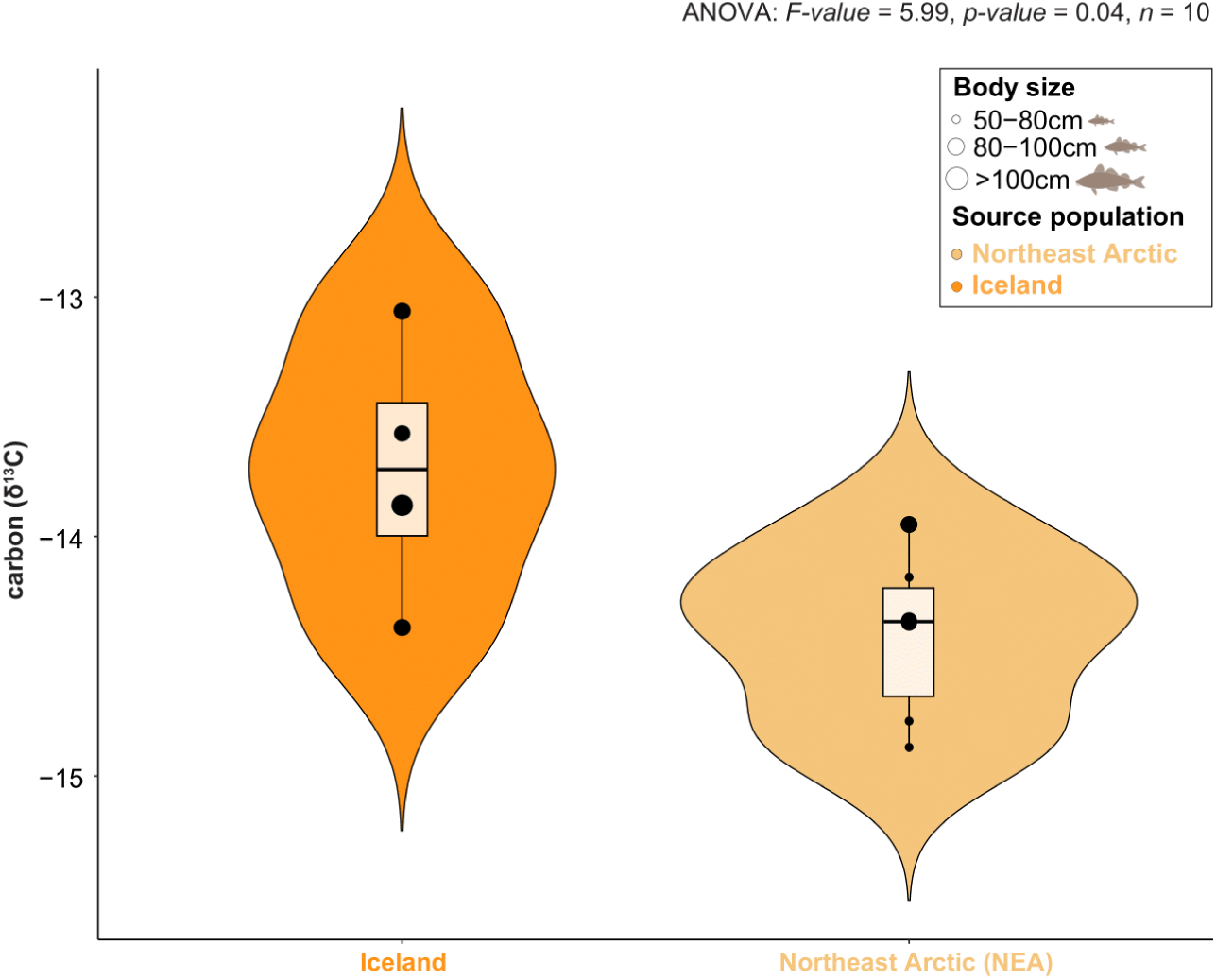
Differences in carbon (δ^13^C) values across specimens with a binary assignment to genetically inferred source populations: Iceland or NEA. Only specimens with a 70%, or higher probability of being assigned to either origin, were used in this analysis (see Table S2, S3).

## 4. Discussion

We have implemented a multidisciplinary approach using aDNA and stable isotopes to identify the biological source and distribution of Atlantic cod specimens from medieval Oslo over a period of 600 years. Combining genomic and isotopic analyses, we found an overall diverse origin of Atlantic cod specimens in medieval Oslo (eleventh to seventeenth centuries). Atlantic cod obtained through trade (e.g., either the Northeast Arctic (NEA) or possibly Iceland) are present throughout time and are found in the earliest deposits. Using multivariate linear regression, we found a significant relation between carbon (δ^13^C) values and the (genomic) probability of being assigned to a NEA biological origin. Moreover, by controlling for Atlantic cod size, PCA analysis supports a separation between the north-central and northernmost genomic groups based on stable isotope values. Our ANOVA results suggest that δ^13^C values are also significantly different between those specimens with different chromosomal inversion LG1 genotypes (NEA and Iceland). Considering that this inversion is associated with different behaviour patterns and habitat preferences in Atlantic cod [46–50], our observation suggests a diverse biological source of Atlantic cod specimens during medieval Oslo. Below we describe the implications of these findings.

Atlantic cod specimens obtained from remote northern fisheries (e.g., NEA or possibly Iceland) are found in Oslo since the eleventh century (*ca.* 1000 CE). This pattern is different from patterns of trade in other locations in Europe, such as in England, Flanders, Poland and Estonia where this long-distance trade increased from the thirteenth to fourteenth centuries onwards [6–8, 61]. S*tockfish* transport from northern Norway to Haithabu, now in northern Germany, has been observed by the eleventh century (i.e., before *ca.* 1066 CE [5]). Thus, our results suggest that trade of Atlantic cod to Oslo probably from the Lofoten or Vesterålen archipelagos appears to be continuous in this region since such earlier periods. As Oslo was not a major hub of *stockfish* trade, it can be inferred that the Atlantic cod of northern origin present since *ca.* 1000 CE were primarily for local consumption.

Following Martínez-García, Ferrari [8], we genetically identified a presumed Icelandic origin of processed fish, possibly *stockfish* with specimens assigned to Icelandic frontal (migratory behavior) or coastal (stationary behavior) ecotypes. Genomic differences between Icelandic and NEA cod can be found in the chromosomal inversion LG1, where a higher frequency of north-central genotypes can be found in Iceland [50]. However, there are similarities between inversion frequencies (for LG1) between deep water Iceland and NEA cod [8, 62]. Considering such similarities and possible biological complexity, the genetic assignments based on inversion frequencies to either Iceland (either frontal and coastal ecotype) or NEA remain uncertain. Nonetheless, NEA cod has distinct migratory behaviour by feeding in the Barents Sea before spawning along the Norwegian coast [63]. The Icelandic populations in contrast, consist of a combination of two ecotypes with different spawning, migratory and feeding behaviour. According to modern otolith increment growth, Icelandic cod grow faster than NEA cod during early stages of life (up to 6 years), whereas NEA cod appears to grow faster during older years [64]. Consequently, significantly higher δ^13^C values in Icelandic cod otoliths (δ^13^C_oto_) compared to NEA have been associated with differences in fish growth and also metabolism [64]. Our genetic assignment results are consistent with such significant difference of δ^13^C values for specimens assigned to NEA based on multivariate linear regression and when compared to those specimens assigned to Iceland; these specimens may have been exposed to different oceanographic and ecological conditions [65, 66] and different metabolic activity [6, 63]. We do not observe overall differences in body size categories between genomic assignments to Iceland and NEA specimens, or northernmost and north-central genomic groups, hence the isotopic differentiation between these genetic categories is not driven by these fish feeding at different trophic level. We do observe significant differences between body size categories across isotopic values (δ^13^C, δ^15^N and δ^2^H), and across time (fish from Brann4 (1275-1375 CE) are larger compared to those from Brann2 (1125-1275 CE) and Brann3 (1250-1325 CE)). These differences between fish of different size reflect the expected ecological complexity during an individual’s lifetime (related to size-specific metabolic rates that decrease as fish grow older), sexual maturation (which differs according to geographical latitude) and diet composition (from lower or higher trophic levels) of different ecotypes or individuals[67–69]. Interestingly, we observed a binomial distribution of δ^13^C and δ^15^N values within the medium size specimens (80-100 cm), which are presumed to feed at similar trophic levels given they are classified in the same size category. These differences might either reflect differences in their environment (perhaps different locations) or distinct feeding strategies amongst individuals of different sizes [70–72]. Overall, our findings, including variability within the stable isotope data, suggest that multiple localities and/or ecotypes, provided *stockfish* to Oslo from the eleventh century onwards.

## Conclusion

This study highlights the utility of combining ancient DNA methods with isotope analysis, to describe the biological origin and biological differences of economically important marine species. For millennia, people have relied on Atlantic cod as a food source and key income product for the development of coastal communities across northern Europe. In fact, our results reveal a continuous presence of Atlantic cod obtained from remote locations like northern Norway or possibly Iceland since the eleventh century in medieval and post-medieval Oslo. This pattern of trade is different from that observed in Britain and Continental Europe outside Scandinavia [8]. Knowledge on the extension of long-distance fish trading patterns can provide valuable information about the exploitation timeline of specific Atlantic cod stocks. While interpretation of the genomic assignment of ancient fish specimens to Iceland remain uncertain based upon inversion frequencies only, the association of genetic data with differences in isotopic values does provide evidence for the existence of mixed fisheries, targeting either fish at different spatial locations, or co-occurring ecotypes that supported the long-distance trade to Oslo.

## Data accessibility

The raw reads for the ancient specimens are released under the ENA accession numbers PRJEB37681 and PRJEB71940.

## Author contributions

B.S and J.H.B. conceived and designed the study. LM-G and AP wrote the original manuscript. AP, LM-G and GF conducted aDNA laboratory work. AH and CK conducted collagen extractions and stable isotope analysis. AP and LM-G performed the genomic analysis. LM-G and AP designed and performed the statistical analyses for isotopic data with input from BS and JHB. Data visualization: LM-G and AP. Fish and bone illustrations: LM-G. AKH, MV, and JHB provided archaeological samples and context. BS and JHB provided funding acquisition. All authors reviewed, discussed and edited the final draft of the manuscript.

## Acknowledgments

We thank J. Rolfe (Cambridge) who assisted with the isotope ratio mass spectrometry, T. O’Connell (Cambridge) who facilitated use of the Dorothy Garrod Laboratory for Isotopic Analysis and Katrien Dierickx for giving advice in statistical analysis. The hydrogen and sulphur isotope analyses were conducted by Iso-Analytical Limited (I. Begley). Finally, we thank M. Skage, S. Kollias and A. Tooming-Klunderud at the Norwegian Sequencing Centre for sequencing and processing of samples. We also thank K.S. Jakobsen and S. Jentoft for insightful discussions and Eric J. Guiry for commenting on an early version of this manuscript. Sequencing analysis were performed on the SAGA Cluster using the resources and assistance from the SIGMA2 Metacenter, the Norwegian National Infrastructure for High Performance Computing and Data Storage.

---

**Figure S1.**
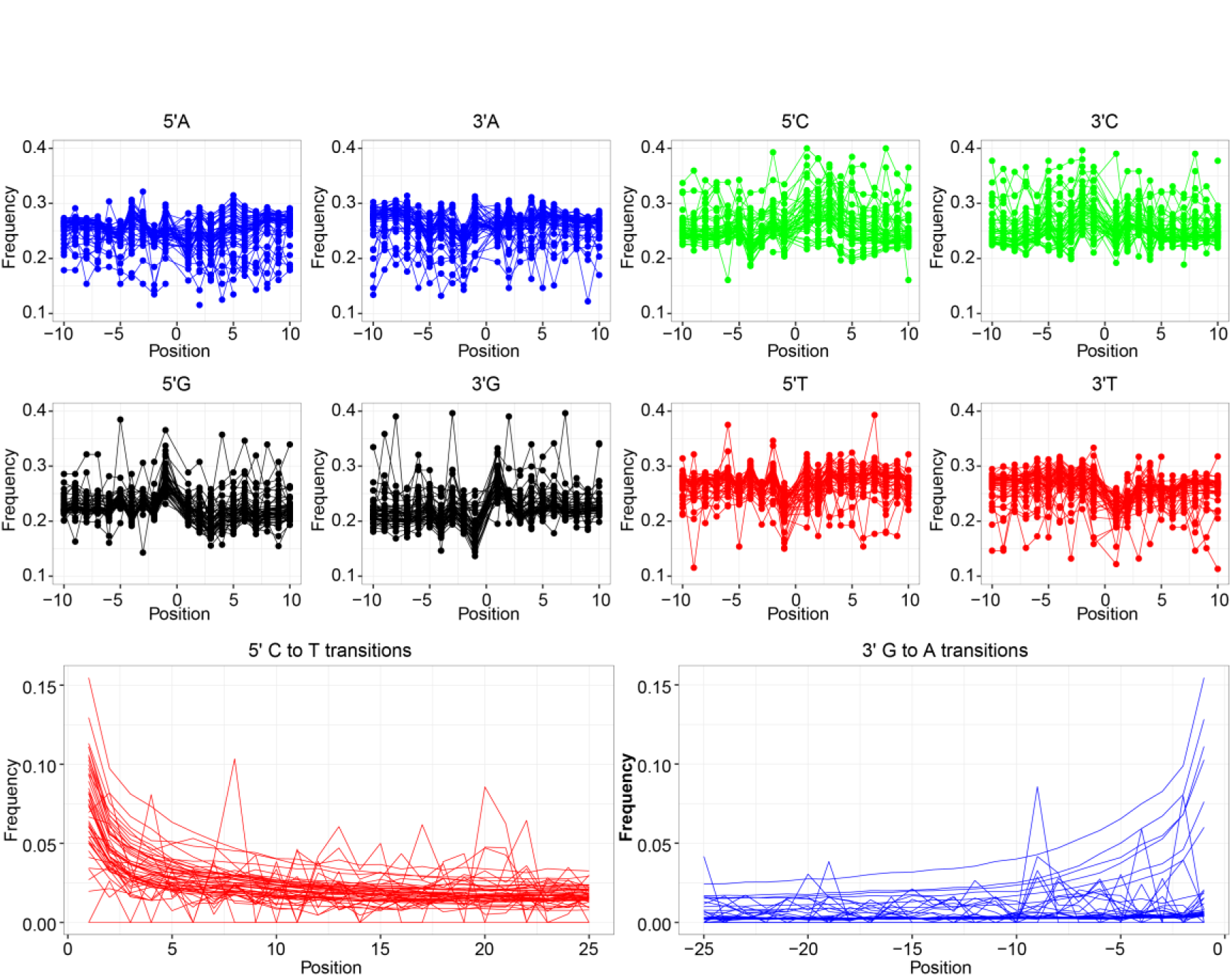
Typical fragmentation and misincorporation patterns of nucleotides of aDNA from sequencing data of 50 Atlantic cod specimens from Oslogate 6 (*n* = 44) and Oslo Mindets tomt (*n* = 6). Base frequencies are shown in the top panel. The bottom panel shows the increase in cytosine to thymine (C > T) misincorporations due to cytosine deamination at the 5′-end of DNA fragments and the corresponding increase of guanine to adenine (G > A) misincorporations at the 3′-end. Several specimens with (lower number of reads <12,000) introduce stochastic patterns in the Cytosine to thymine (C > T & G > A) misincorporations.

**Figure S2.**
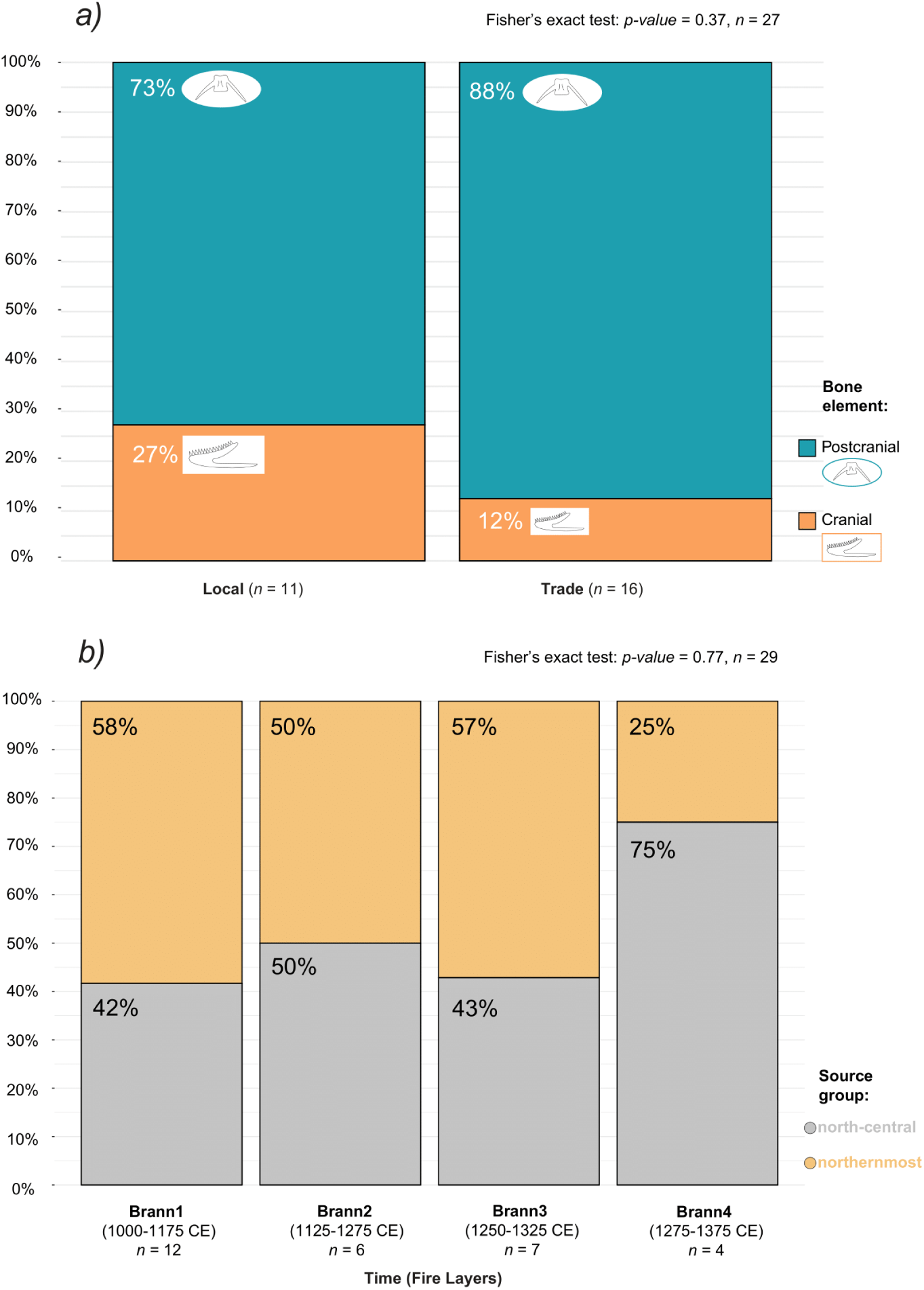
***(a)*** Association between bone element (cranial or postcranial) and specimens with a presumed local or traded origin (*p-value* = 0.37). Percentage (%) of cranial (orange) or postcranial (blue) bones with a local or traded (NEA or Iceland) genomic assignment (*n* = 27). Individuals with ambiguous origin (i.e., COD082, COD084, COD339, COD356, COD398, COD405) and individuals with a northernmost origin below 70% probability (i.e., COD408 and COD421) were excluded from this test. ***(b)*** Distribution of specimens genetically assigned to a binary origin to the north-central (grey) or the northernmost (orange) groups across time (Brann 1 to Brann 4; *p-value* = 0.77, *n* = 29).

**Figure S3.**
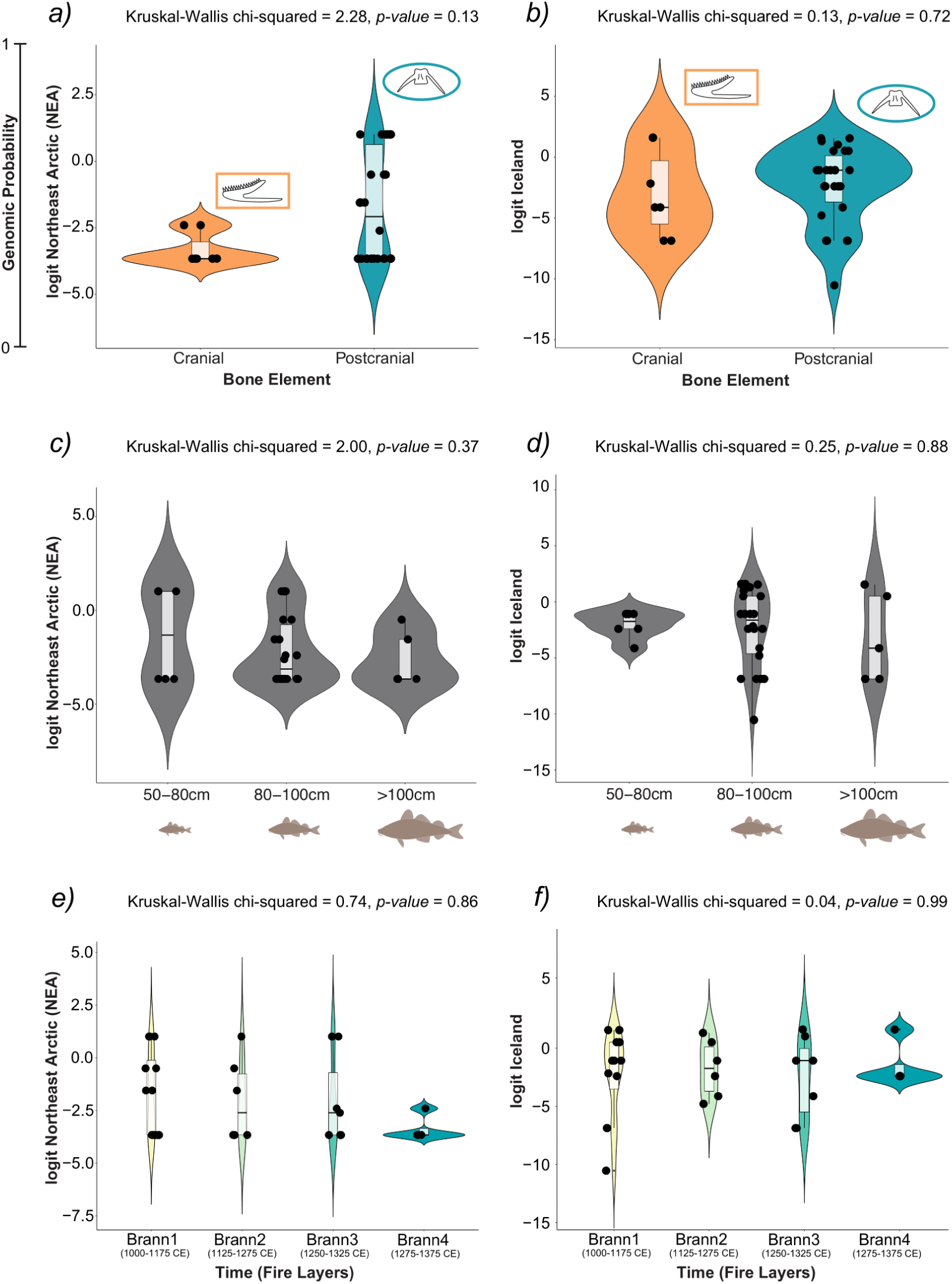
Associations between the probability of having a Northeast Arctic (NEA) or Icelandic origin (*logit* values) and ***(a)*** bone element (cranial or postcranial) categories, ***(b)*** size categories (small: 50-80 cm, medium: 80-100 cm, large: >100 cm), and ***(c)*** across time (Brann1 to Brann4).

**Figure S4.**
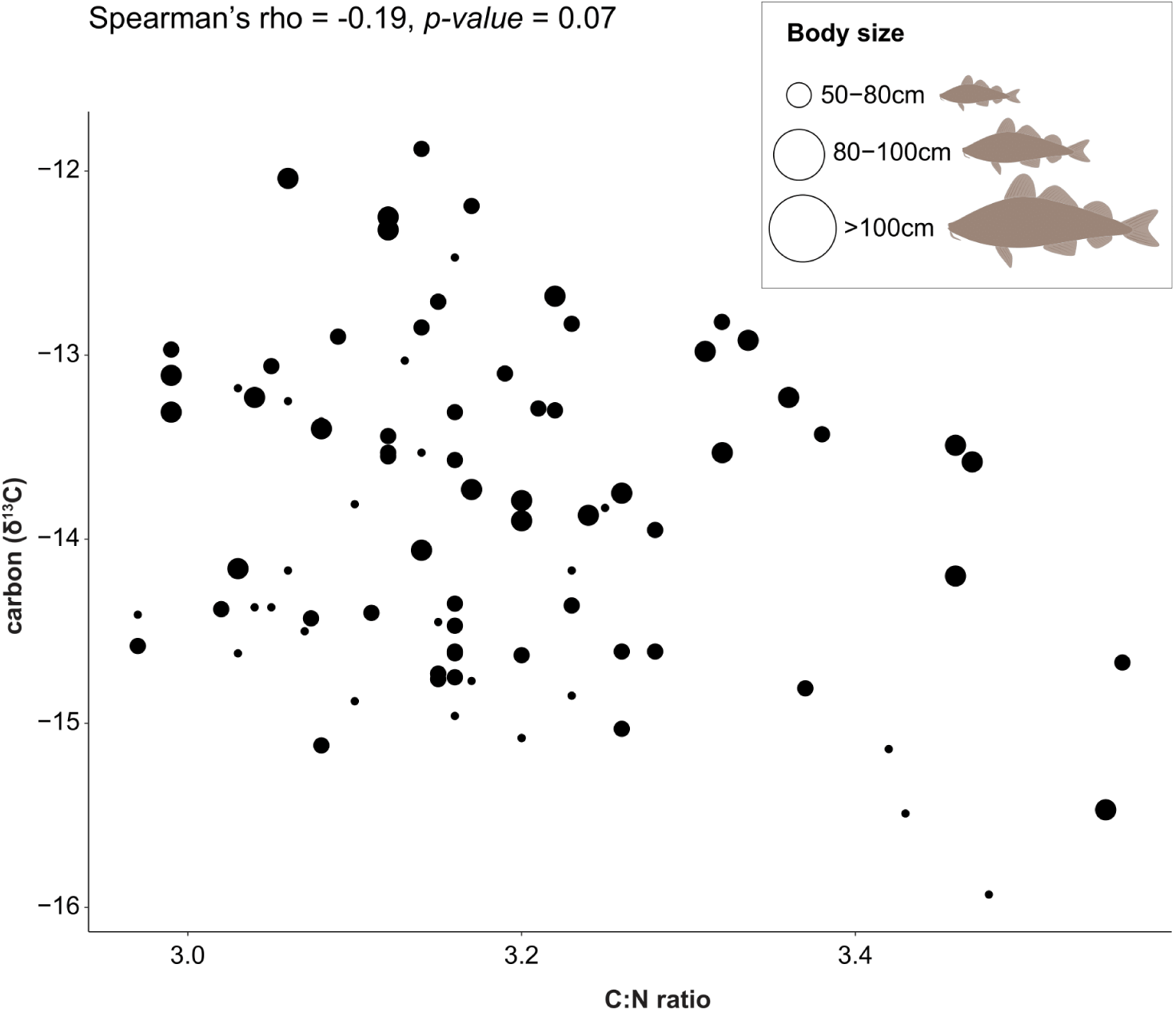
Isotope correlation between carbon (δ^13^C) values and C:N ratios from archaeological Atlantic cod bones from medieval Oslo. A non-significant correlation is observed.

**Figure S5.**
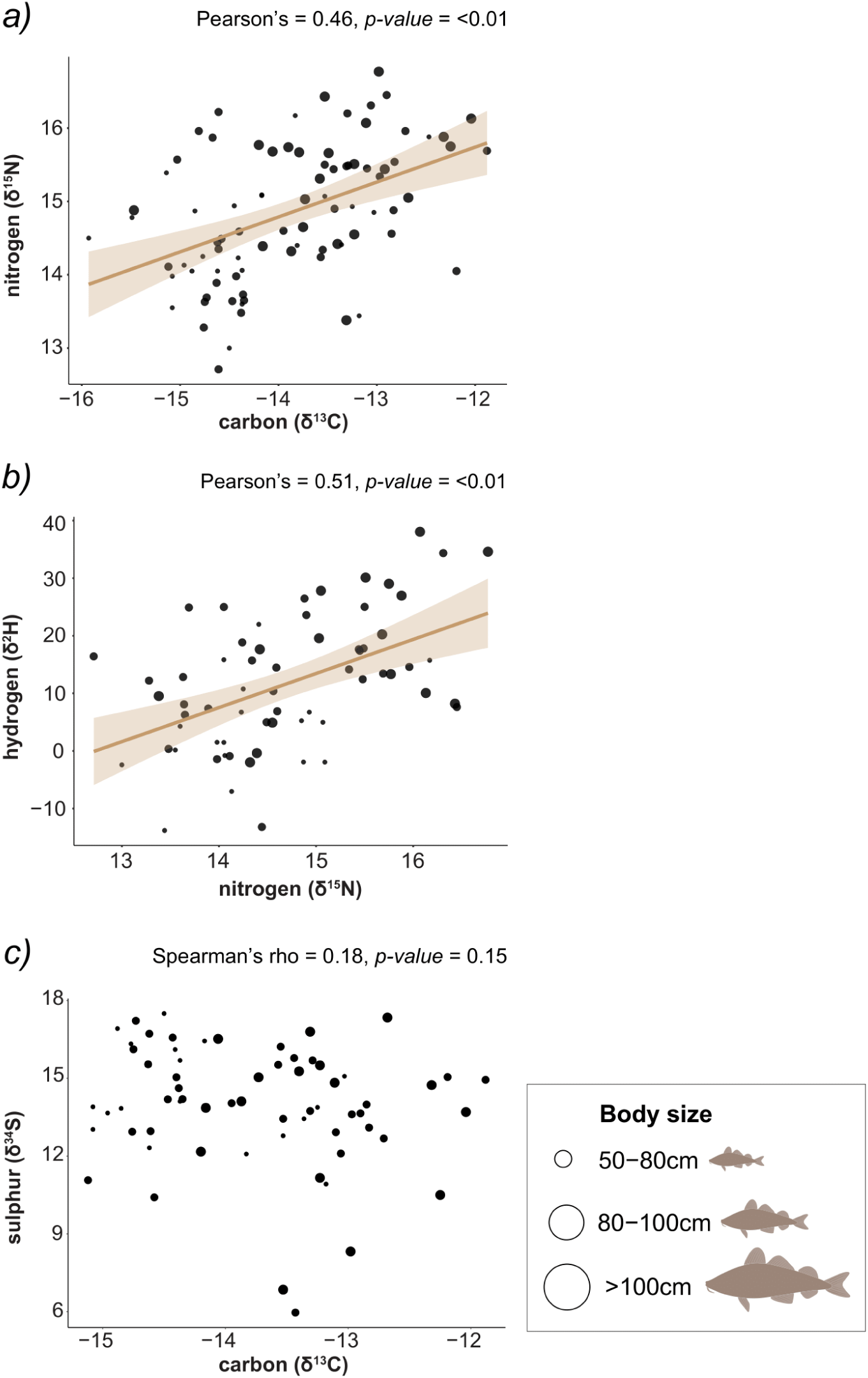
Isotope correlations of archaeological Atlantic cod bones from medieval Oslo. ***(a)*** Positive correlations between carbon and nitrogen (δ^13^C-δ^15^N), and ***(b)*** nitrogen and hydrogen (δ^2^H) isotopes. ***(c)*** Negative correlation between sulphur and carbon (δ^34^S-δ^13^C).

**Figure S6.**
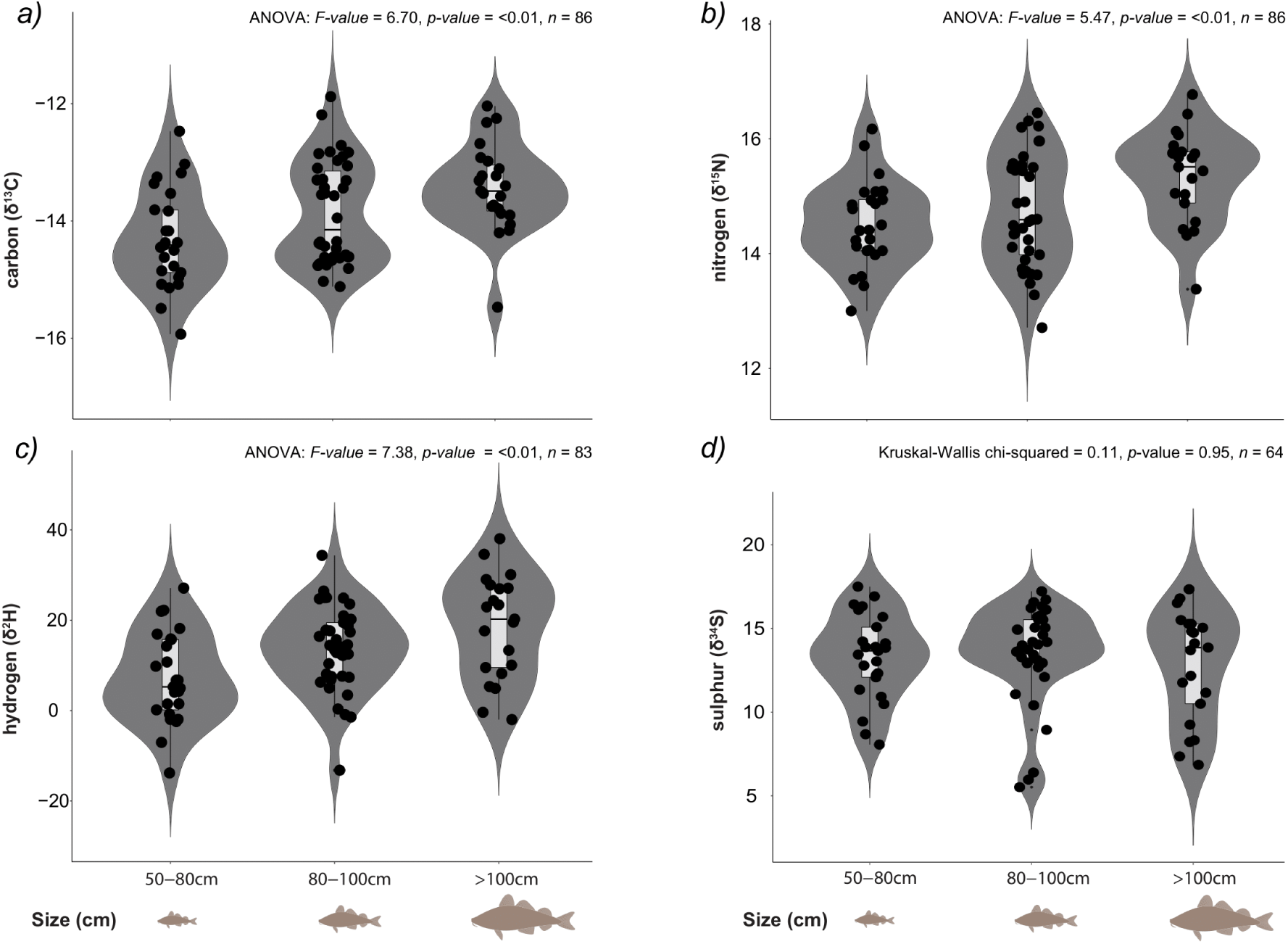
Differences between isotope values across body size and ***(a)*** carbon (δ^13^C) values, ***(b)*** nitrogen (δ^15^N) values, ***(c)*** hydrogen (δ^2^H) values, and ***(d)*** sulphur (δ^34^S) values. Significant differences were found in δ^13^C, δ^15^N and δ^2^H values between the smaller (50-80 cm) compared to the larger (>100 cm) body size categories (TukeyHSD post-hoc test *p-value* = <0.01). Bimodal distributions were observed within carbon (δ^13^C) and nitrogen (δ^15^N) values for the medium size category of fish (see also Figure S7.)

**Figure S7.**
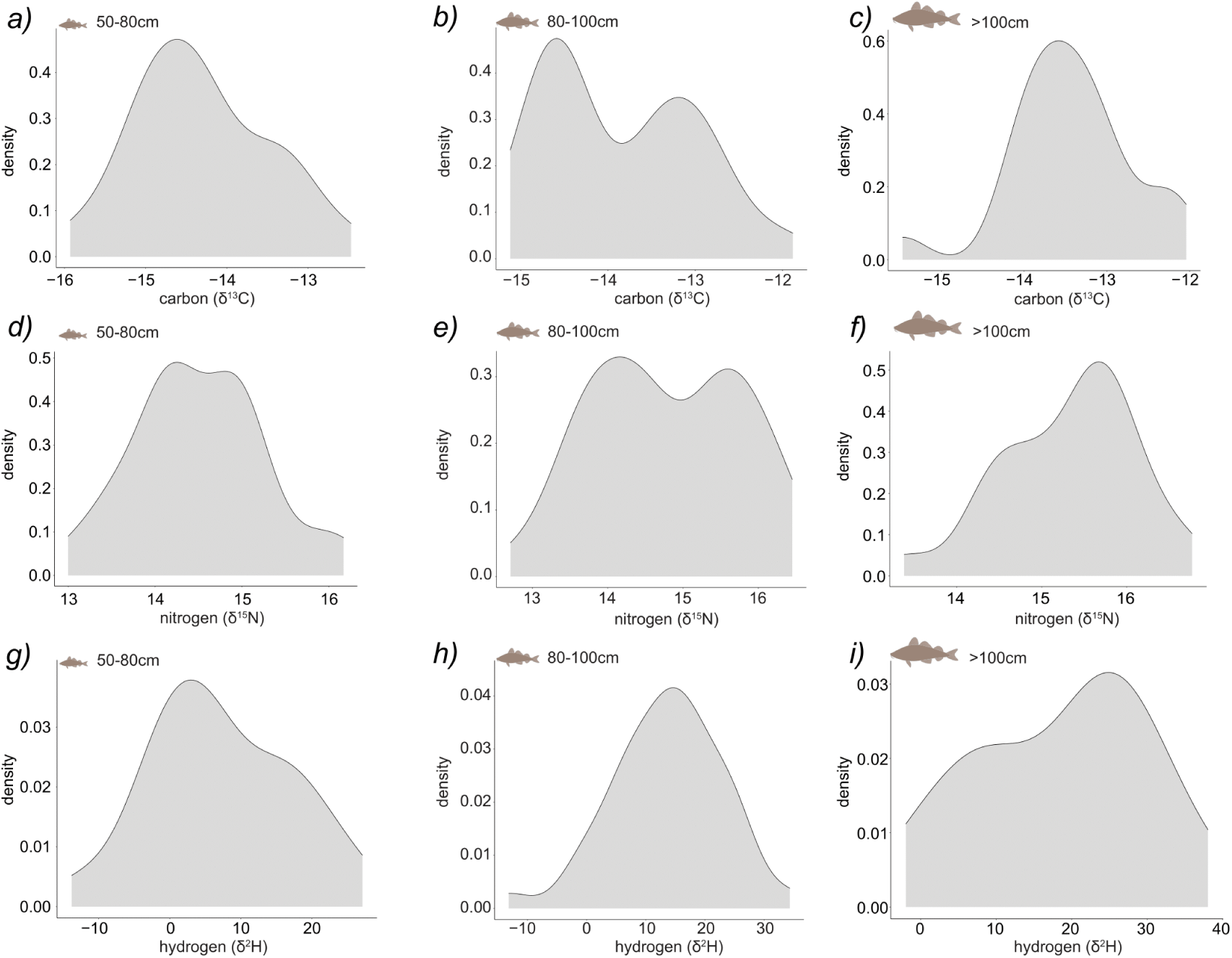
Distribution of ***(a, b, c)*** carbon (δ^13^C), ***(d, e, f)*** nitrogen (δ^15^N), and ***(g, h, i)*** hydrogen (δ^2^H) values within body size categories (small: 50-80 cm, medium: 80-100 cm, large: >100 cm). A bimodal distribution is observed only within medium sized fish (80-100cm) for δ^13^C and δ^15^N but not in δ^2^H or smaller and larger body size categories.

**Figure S8.**
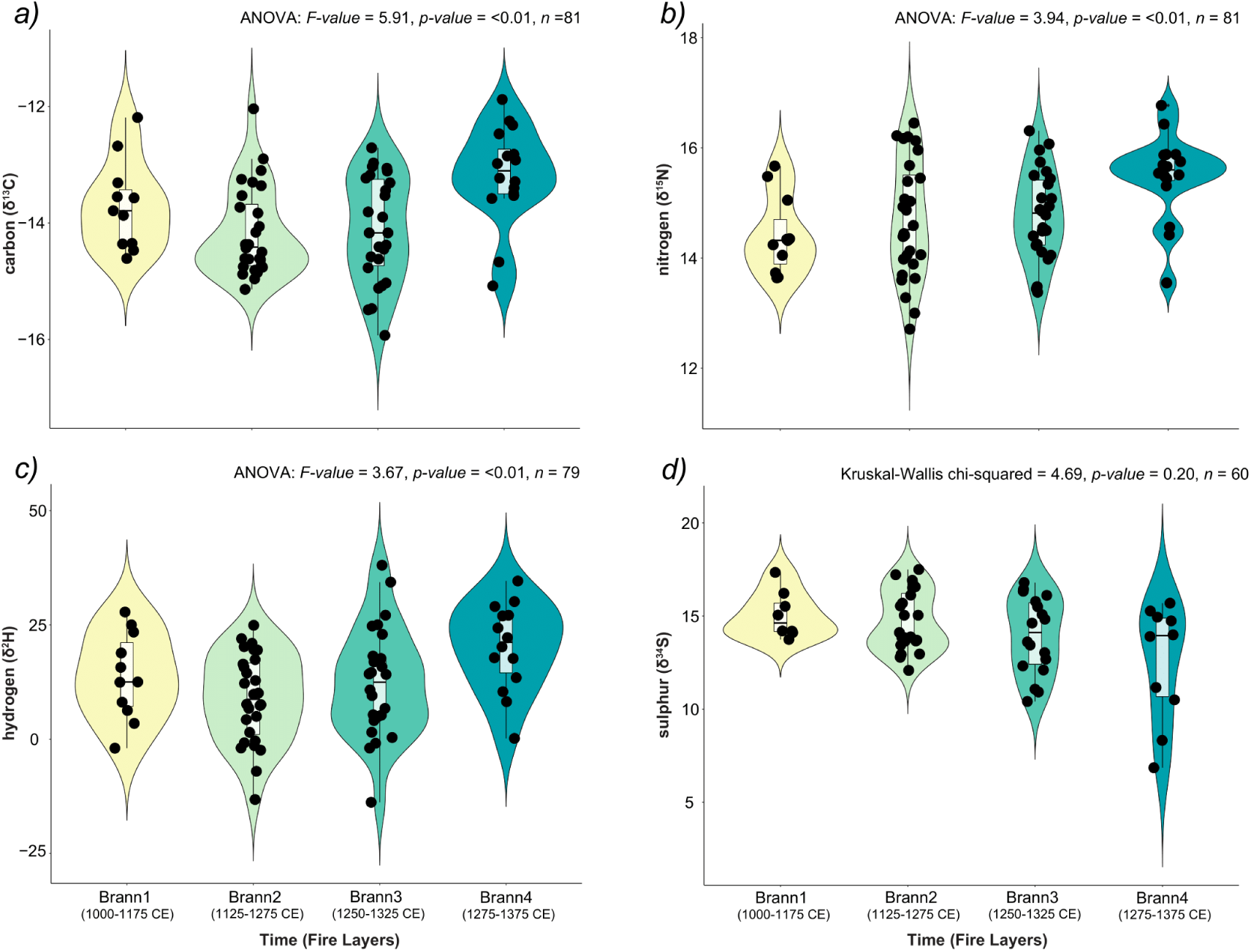
Differences between isotope values across time (Brann1 to Brann4): ***(a)*** carbon (δ^13^C) values, ***(b)*** nitrogen (δ^15^N) values, ***(c)*** hydrogen (δ^2^H) values, and ***(d)*** sulphur (δ^34^S) values. Significant higher isotope values (δ^13^C *p-value* = <0.01; δ^15^N *p-value* = 0.03 and δ^2^H *p-value* = 0.01) were found between Brann4 (1275-1375 CE) and Brann2 (1125-1275 CE). Furthermore, significant higher δ^13^C values (*p-value* = <0.01) were found between Brann4 compared to Brann3 (1250-1325 CE), and significantly higher δ^15^N values (*p-value* = 0.01) were found between Brann4 and Brann1 (1000-1175 CE). No significant correlations were obtained between δ^34^S values across time (Kruskal-Wallis chi-squared = 4.69, df = 3, *p-value* = 0.20, *n* = 60).

**Figure S9.**
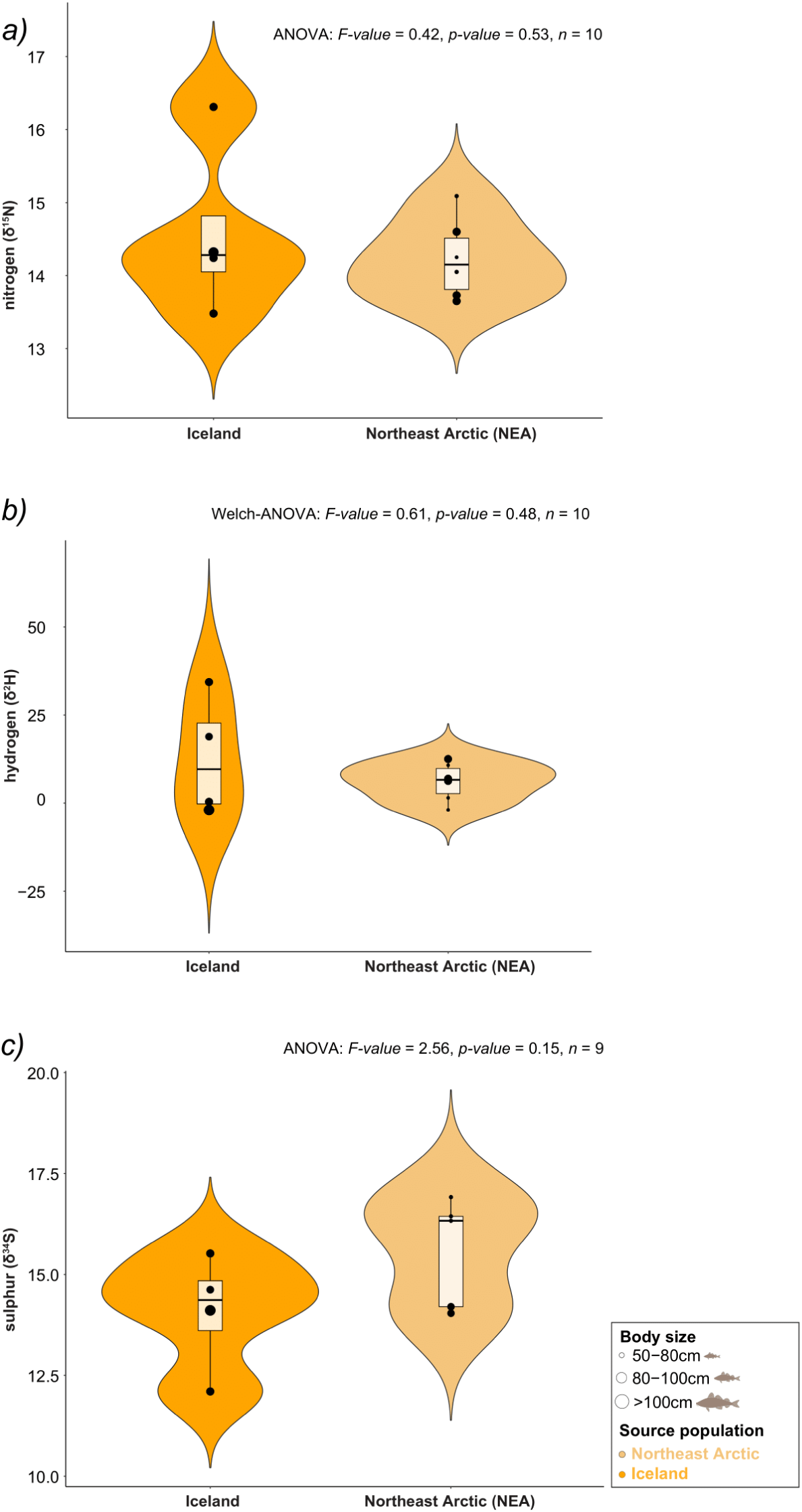
Differences across a binary assignment to Iceland or NEA populations and ***(a)*** nitrogen (δ^15^N) values, ***(b)*** hydrogen (δ^2^H) values, and ***(c)*** sulphur (δ^34^S) values. Only specimens with a 70%, or higher probability of being assigned to either origin, were used.

